# Evaluating *aroA* gene essentiality and EPSP synthase vulnerability in *Mycobacterium smegmatis* under different nutritional conditions

**DOI:** 10.1101/2020.03.02.974360

**Authors:** Mario Alejandro Duque-Villegas, Bruno Lopes Abbadi, Paulo Ricardo Romero, Luiza Galina, Pedro Ferrari Dalberto, Valnês da Silva Rodrigues-Junior, Candida Deves Roth, Raoní Scheibler Rambo, Eduardo Vieira de Souza, Marcia Alberton Perello, Pablo Machado, Luiz Augusto Basso, Cristiano Valim Bizarro

## Abstract

The epidemiological importance of bacteria from the genus *Mycobacterium* is indisputable and the necessity to find new molecules that can inhibit their growth is urgent. The shikimate pathway, required for the synthesis of important metabolites in bacteria, represents a target for inhibitors of *Mycobacterium tuberculosis* growth. The *aroA*-encoded 5-enolpyruvylshikimate-3-phosphate synthase (EPSPS) enzyme catalyzes the sixth step of the shikimate pathway. In this study, we combined gene knockout, gene knockdown and kinetic assays to evaluate *aroA* gene essentiality and the vulnerability of its protein product, EPSPS synthase from *Mycobacterium smegmatis* (*Ms*EPSPS), under different nutritional conditions. We demonstrate by an allelic exchange-based gene knockout approach the essentiality of *Ms*EPSPS under rich and poor nutritional conditions. By performing gene complementation experiments with wild-type (WT) and point mutant versions of *aroA* gene, together with kinetic assays using WT and mutant recombinant proteins, we show that *aroA* gene essentiality depends on *Ms*EPSPS activity. To evaluate *Ms*EPSPS vulnerability, we performed gene knockdown experiments using the Clustered Regularly Interspaced Short Palindromic Repeats interference (CRISPRi) system. The experiments were performed in both rich and defined (poor) media, using three different repression forces for *aroA* gene. We only observed growth impairment when bacteria were grown in defined medium without supplementation of aromatic amino acids, thereby indicating that *Ms*EPSPS vulnerability depends on the environment conditions.

**Importance:** We evaluated both gene essentiality and target vulnerability of the enzyme that catalyzes the sixth step of the shikimate pathway, the *aroA*-encoded 5-enolpyruvylshikimate-3-phosphate synthase from *Mycobacterium smegmatis* (MsEPSPS). Combining gene knockout experiments and kinetic assays, we established a causal link between *aroA* gene essentiality and the biological function of EPSPS protein, which we advocate is an indispensable step for target validation. Moreover, we characterized *Ms*EPSPS vulnerability under different nutritional conditions and found it is a vulnerable target only when *M. smegmatis* is grown under poor nutritional conditions without supplementation with aromatic amino acids. Based on our findings, we suggest that gene essentiality information should be obtained from gene knockout experiments and not knockdown approaches, as even low levels of a protein after gene silencing can lead to a different growth phenotype when compared to that under its complete absence, as was the case with *aroA* and *Ms*EPSPS in our study.

## INTRODUCTION

Human tuberculosis (TB) is an important infectious disease that continues to be a public health threat worldwide. Despite the joint global efforts to lower TB burden, which resulted in a 6.3% and 11% cumulative decline in incidence and mortality, respectively, between 2015 and 2018, the End TB Strategy milestones for 2020 are far from being reached [1]. According to the last World Health Organization (WHO) TB report, around 10.0 million people developed the disease, and 1.2 million HIV-negative individuals died from it in 2018 [1]. In humans, the acid-fast *Mycobacterium tuberculosis* bacilli is the main causative agent of pulmonary TB, a debilitating condition that is fatal without treatment [1, 2]. Although TB is considered a curable disease, with a success rate of approximately 85% for drug-susceptible strains, the spread of multidrug-resistant and rifampicin-resistant TB (MDR/RR-TB) poses a challenge to the current first-line treatment [1]. Resistance cases of TB have been documented since the very beginning use of streptomycin as the first anti-TB monotherapy in 1943 [3, 4]. Therapies for MDR/RR-TB are complex, more expensive, longer, and more toxic, when compared to those for drug-susceptible TB, and it is estimated that only 56% of MDR-TB cases reach cure [1, 5].

The spread of resistant strains is related to the acquisition of multiple molecular mechanisms that allows *M. tuberculosis* evade the action of anti-TB drugs, mostly by mutations on drug targets [6]. Therefore, the development of new anti-TB drugs having new mechanisms of action are needed. As the number of molecular targets of current bacterial agents are limited [7], there is a growing interest in finding and validating new molecular targets for drug development. Apart from being essential, a drug target should be vulnerable, which means that the incomplete inhibition of its activity is sufficient to produce a lethal phenotype [8]. Some genes/proteins were found to be essential but not vulnerable, prompting the need to use molecular genetic tools to study target vulnerability as part of the target validation process [9, 10].

The shikimate pathway is considered an attractive target for the development of rational-based new antimicrobial agents. It is essential for the growth of bacteria, but absent in most animals, including mammals, favoring the development of selective inhibitors for pathogenic bacteria [11, 12]. This pathway is composed by seven different enzymatic steps, leading to the production of chorismate, which is a precursor of naphthoquinones, menaquinones and mycobactins as well as folates, ubiquinones, tryptophan, tyrosine and phenylalanine [11,13,14].

The 5-enolpyruvylshikimate-3-phosphate synthase (EPSPS; EC 2.5.1.19) is the sixth enzyme of the shikimate pathway. EPSPS is coded by *aroA* gene and catalyzes the transfer of the carboxyvinyl portion of the phosphoenolpyruvate substrate (PEP) to the carbon-5 hydroxyl group of shikimate 3-phosphate (S3P) to form the enolpyruvyl shikimate-3-phosphate product (EPSP) [15]. The *aroA* gene was found to be essential for growth or virulence for many bacterial species, such as *Streptococcus pneumoniae*, *Bordetella bronchiseptica*, *Salmonella typhimurium* and species of the genus *Aeromonas* and *Shigella* [16, 17]. However, the vulnerability of EPSPS as a drug target was not experimentally studied yet.

In this study, we used *Mycobacterium smegmatis* as a model organism to evaluate the essentiality of *aroA* gene and the vulnerability of its protein product (*Ms*EPSPS). The *aroA* gene was knocked out in allelic exchange experiments and found to be essential for *M. smegmatis* growth under the conditions tested. We also evaluated the ability of a wild-type (WT) *aroA* gene or *aroA* alleles containing point mutations to complement the knockout (KO) *aroA* strain. Specifically, we found two EPSPS amino acid residues as essential (R134 and E321). Mutated versions of recombinant *Ms*EPSPS were expressed, purified and their kinetic activities characterized. We found that recombinant R134 and E321 mutants have diminished EPSPS enzyme activity. Our results suggest that the *aroA* essentiality under our experimental conditions depends on the EPSPS activity of its protein product. Moreover, using the CRISPRi system [18], we evaluated EPSPS vulnerability under different nutritional conditions. Interestingly, we found EPSPS as a vulnerable target only when grown on 7H9 medium. When supplemented with aromatic amino acids (7H9 + L-tryptophan + L-phenylalanine + L-tyrosine) or grown on a rich medium (LB), we observed normal bacterial growth of CRISPRi-inactivated *aroA* gene.

## MATERIALS AND METHODS

### Bacterial Strains, Growth Conditions and Transformation

*Escherichia coli* DH10B strain was used for all cloning procedures and routinely grown in LB medium (broth and agar), at 37°C. *Mycobacterium smegmatis* mc²155 strain [19] was kindly provided by Dr. William R. Jacobs, Jr., Albert Einstein College of Medicine, NY, USA. *M. smegmatis* was used for gene knockout and knockdown experiments, and it was grown in LB medium, Difco™ Middlebrook 7H9 broth (Becton Dickinson – BD), supplemented with 0.05% (v/v) Tween 80 (Sigma-Aldrich), and 0.2% (v/v) glycerol (MERCK), or Difco™ Middlebrook 7H10 Agar (BD), supplemented with 0.5% (v/v) glycerol. Wherever required, the following antibiotics or small molecules were used: 50 µg/mL ampicillin (Amp - Sigma-Aldrich), 25 µg/mL kanamycin (Kan - Sigma-Aldrich) for culturing recombinant *E. coli* strains. Also, 25 µg/mL Kan, 50 µg/mL hygromycin (Hyg – Invitrogen), 100 ng/mL anhydrotetracycline (ATC – Sigma-Aldrich), 50 µg/mL of each amino acids L-tryptophan (FisherBiotech), L-phenylalanine (Sigma-Aldrich) and L-tyrosine (Sigma-Aldrich) for culturing *M. smegmatis* strains. All *E. coli* strains were routinely transformed by electroporation using cuvettes of 0.2 cm, with a 200 Ω resistance, 25 µF capacitance, pulse of 2.25 kV for 3 seconds. On the other hand, for *M. smegmatis* strains, the resistance was changed to 1000 Ω and the pulse to 2.5 kV also for 3 seconds [20]. All primers used in this study are listed in Table 1.

**Table 1.**
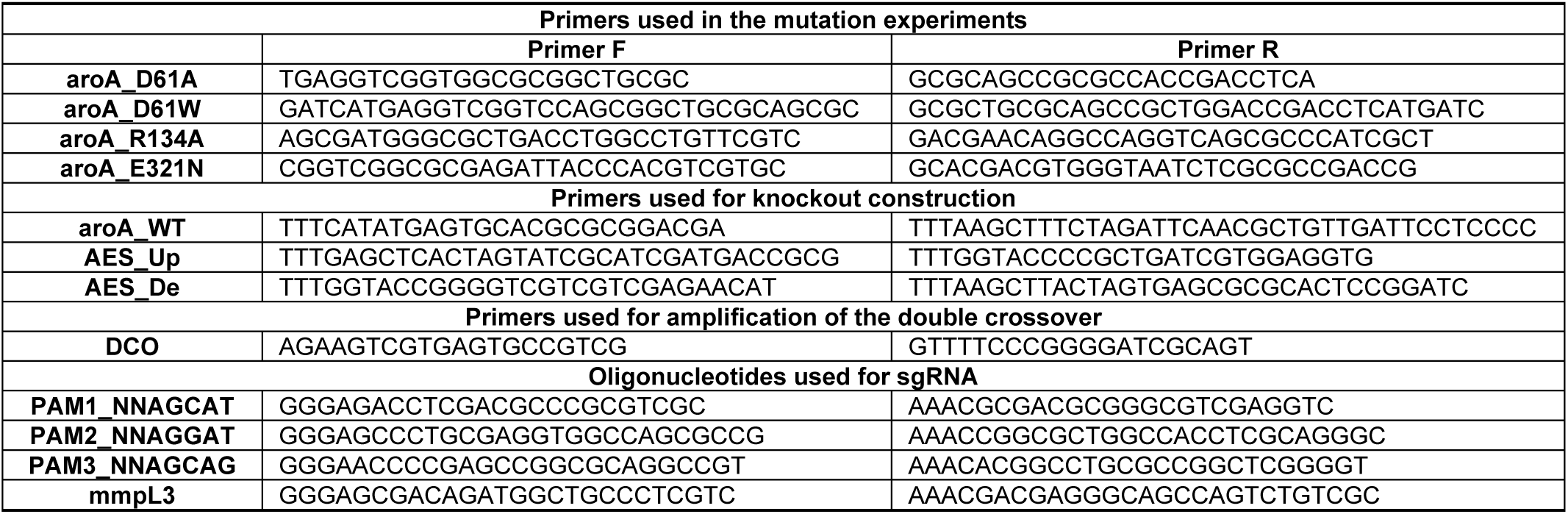
List of oligonucleotides and primers used in this study.

### Construction of vectors for recombinant protein expression

The WT *aroA* gene (MSMEG_1890), predicted to code for a 5-enolpyruvylshikimate-3-phosphate synthase (EPSPS), was amplified by PCR using aroA_WT_Primer F and aroA_WT_Primer R (Table 1), 25 ng of genomic DNA of *M. smegmatis* and 10% of DMSO. Genomic DNA was extracted and purified according to a published protocol [21]. The PCR product of 1,354 bp was gel purified, cloned into the pCR™-Blunt (ThermoFisher) vector, and subcloned into the pET-23a(+) (Novagen) expression vector, using NdeI and HindIII restriction sites. Besides, the pET-23a(+)::*aroA*(WT) recombinant vector was used as template for mutagenesis reactions. Four different mutations (D61A, D61W, R134A and E321N) were incorporated into the gene sequence using the QuickChange XL site-directed mutagenesis kit (Stratagene), along with a set of several primers: primers F and R of aroA_D61A, aroA_D61W, aroA_R134A and aroA_E321N (Table 1). Recombinant clones were confirmed by DNA sequencing.

### Construction of the gene knockout (KO) vector

The genomic flanking sequences of *aroA* gene from *M. smegmatis* were PCR-amplified to serve as allelic exchange substrates (AESs) for gene KO. The upstream flanking sequence (1,066 bp), named AES_Up, included 164 bp of the 5’-end of a*roA*, and it was amplified using AES_Up_Primer F and AES_Up_Primer R (Table 1), containing SacI/Spel and KpnI restriction sites, respectively. The downstream flanking sequence (1,043 bp), named AES_Dw, included 109 bp of the 3’-end of *aroA*, and it was amplified using AES_Dw_Primer F and AES_Dw_Primer R, containing KpnI, and SpeI/HindIII, respectively (Table 1). The AES_Up sequence was cloned into the pUC19 vector, using restriction sites for SacI and KpnI, followed by the AES_Dw insertion using KpnI and HindIII restriction sites. Both AES sequences were confirmed by DNA sequencing. Next, the vector was digested with KpnI, the cohesive endings were filled with *Pfu* DNA polymerase (QuatroG P&D), and the resulting vector was dephosphorylated by CIP (Invitrogen). Then, a 1.2 kb kanamycin resistance cassette from the pUC4K vector was ligated between the AESs. Finally, the whole construction was cut out from pUC19 with SpeI, and inserted into the SpeI site of pPR27xylE vector [22], to yield plasmid pPR27::KO_aroA (Table 1), used to perform allelic replacement.

### Construction of gene complementation (CO) vectors

The wild-type (WT) and the four mutant *aroA* sequences (D61A, D61W, R134A and E321N) were transferred from pET-23a(+) vector to the pMVHG1 shuttle vector [23], using the NdeI and HindIII restriction sites. Each gene sequence (WT and mutants) was ligated downstream to the heat shock promoter P*_hsp60_*. Then, the P*_hsp60_*::*aroA* sequences were cut out with XbaI, gel-purified, and inserted into the XbaI-dephosphorylated site of the pNIP40/b plasmid, yielding five different rescue plasmids (Table 1).

### Construction of gene knockdown (KD) vectors

The vulnerability of *aroA* gene was evaluated using the CRISPRi system, developed by Rock and colleagues [18]. The PLJR962 vector backbone was linearized by BsmBI digestion and gel-purified. Three small-guide RNAs (sgRNAs) scaffolds were built to target different regions of the *aroA* coding sequence (Fig. 1). They were designed to bind the non-template (NT) strand of *aroA* gene, in regions where three different PAM (protospacer adjacent motif) sequences (Table 1) were identified using an in-house script written in Python and made publicly available in the GitHub repository (https://github.com/Eduardo-vsouza/sgRNA_predictor). A sgRNA targeting the *mmpL3* (MSMEG_0250) gene was used as a positive control of knockdown experiments.

**Figure 1.**
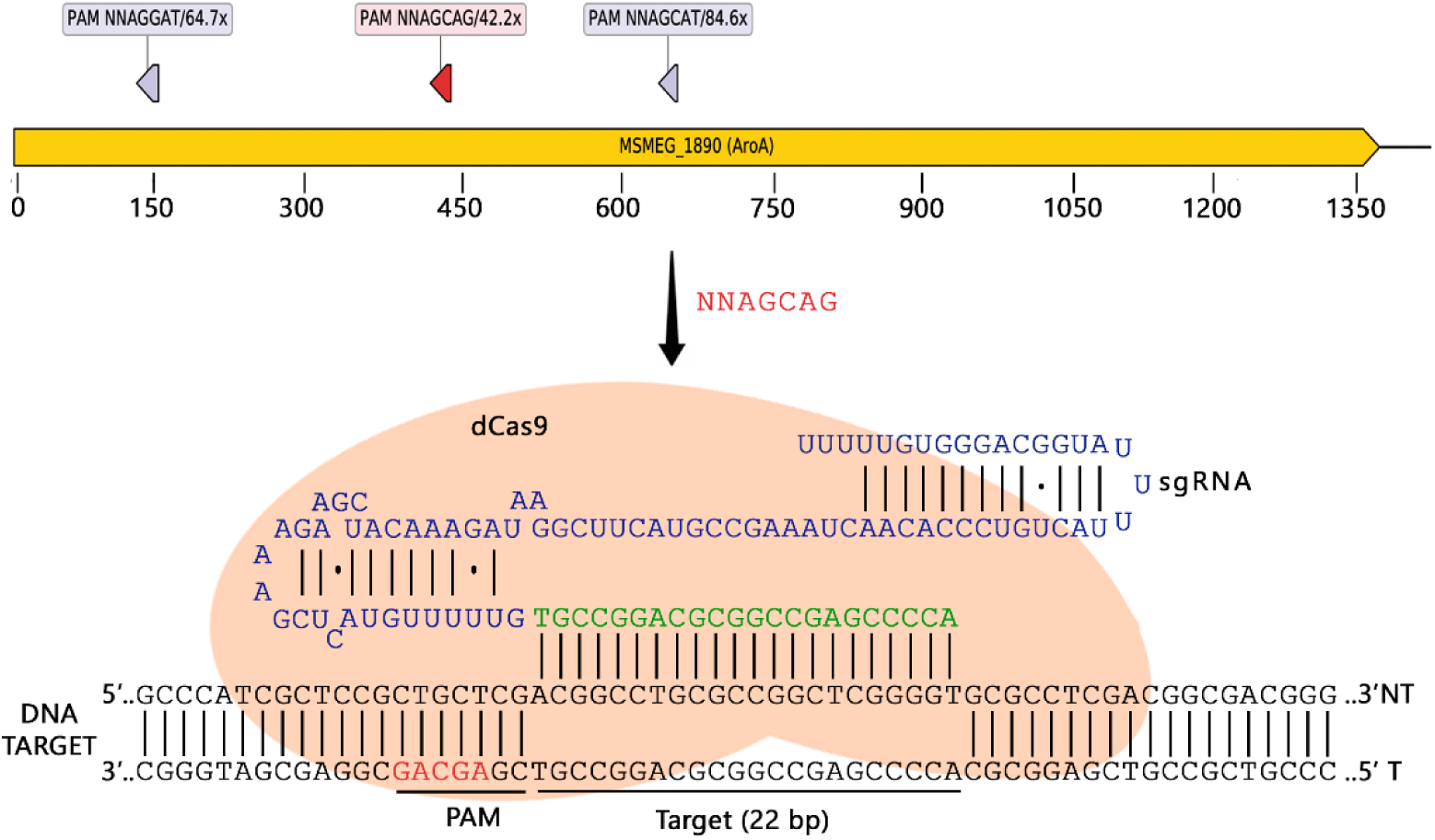
Knockdown of EPSPS-encoding gene *aroA* in *M. smegmatis* using CRISPR interference (CRISPRi). **Upper part:** Location of PAM sequences inside *aroA* locus used in this study. From left to right: 5’-NNAGGAT-3’, 5’-NNAGCAG-3’ and 5’-NNAGCAT-3’. The repression strength of each PAM sequence, according to Rock et al. (2017), is also depicted. **Lower part**: Schematic representation of CRISPRi system associated with *aroA* locus at a target region adjacent to PAM “5’-NNAGCAG-3’”. Dead Cas9 (dCas9) is represented in peach color, sgRNA as a single RNA chain in blue with annealing portion in green and paired with the non-template (NT) strand of target DNA. The “5’-AGCAG-3’” from PAM is depicted in red in the template (T) strand of target DNA in 3’-5’ orientation.

Two partially complementary oligonucleotides (20-25-nt in length) were designed for each sgRNA scaffold (PAM1, PAM2 and PAM3 primers F and R - Table 1). The first nucleotide of each sgRNA started with an “A” or “G” to ensure high transcription efficiency. After annealing (95°C for 5 min; decrease 0.1°C/sec until reaching 25°C), oligos retain single-strand 5’-ends that are complementary to the cohesive ends of BsmBI-digested CRISPRi vector. The ligation of sgRNA scaffolds into the vector backbone using T4 DNA ligase (23°C for 16 h) was confirmed by BsmBI digestion and DNA sequencing.

### Knockout of the *aroA* gene from *M. smegmatis* and gene complementation

The strategy used for gene knockout experiments was based on a published protocol [24]. First, all rescue plasmids (100-300 ng) were introduced into thawed electrocompetent *M. smegmatis* cells (200 µL) by electroporaton, including a control empty vector (pNIP40::Ø), and selected for their hygromycin resistance (Hyg^R^) at 37°C. After three incubation days, one isolated colony of each transformation was grown in 5 mL of LB medium, and electrocompetent cells were prepared again. Then, the pPR27::KO_*aroA* vector was introduced into each merodiploid strain, and transformants were selected on solid LB + kanamycin + hygromycin, at 32°C (permissive temperature), for their kanamycin resistance (Kan^R^), and for the presence of the *xylE* reporter gene (*xylE*^+^) with the addition of a drop of 1% catechol solution (Sigma-Aldrich). Three yellow colonies (Kan^R^, Hyg^R^, XylE^+^) were grown in liquid LB, with kanamycin and hygromycin, at 32°C and 180 rpm, until reaching an optical density (OD) of 0.6-1.0 at 600 nm. Approximately 1 x 10^7^ CFUs were then plated on solid LB + kanamycin + hygromycin + 2% sucrose counter-selective plates, in triplicate, and incubated at 39°C for five days. The inoculum was determined by plating each culture on the same medium, but in the absence of sucrose, at the permissive temperature (32°C), for seven days. Plates were analyzed for the presence of recombinant white colonies (Kan^R^, Hyg^R^, XylE^-^, Suc^R^) with a drop of catechol. Where possible, white colonies were selected to have their genomic DNA extracted to unambiguously confirm the allelic exchange event in the *aroA* locus by PCR. Amplification reactions were performed with DCO_Primer F and DCO_Primer R (Table 1). DCO_Primer F anneals upstream from *aroA* locus, outside the recombination region, while DCO_Primer R anneals inside the kanamycin resistance cassette (Fig. 2A). An amplicon of 1,813 bp in length was obtained from allelic exchange mutants that underwent a double crossover event (Fig. 2B). The WT genomic DNA of WT *M. smegmatis* was used as a negative control. The same experiment was performed in 7H9 and 7H10 broths for comparison purposes.

**Figure 2.**
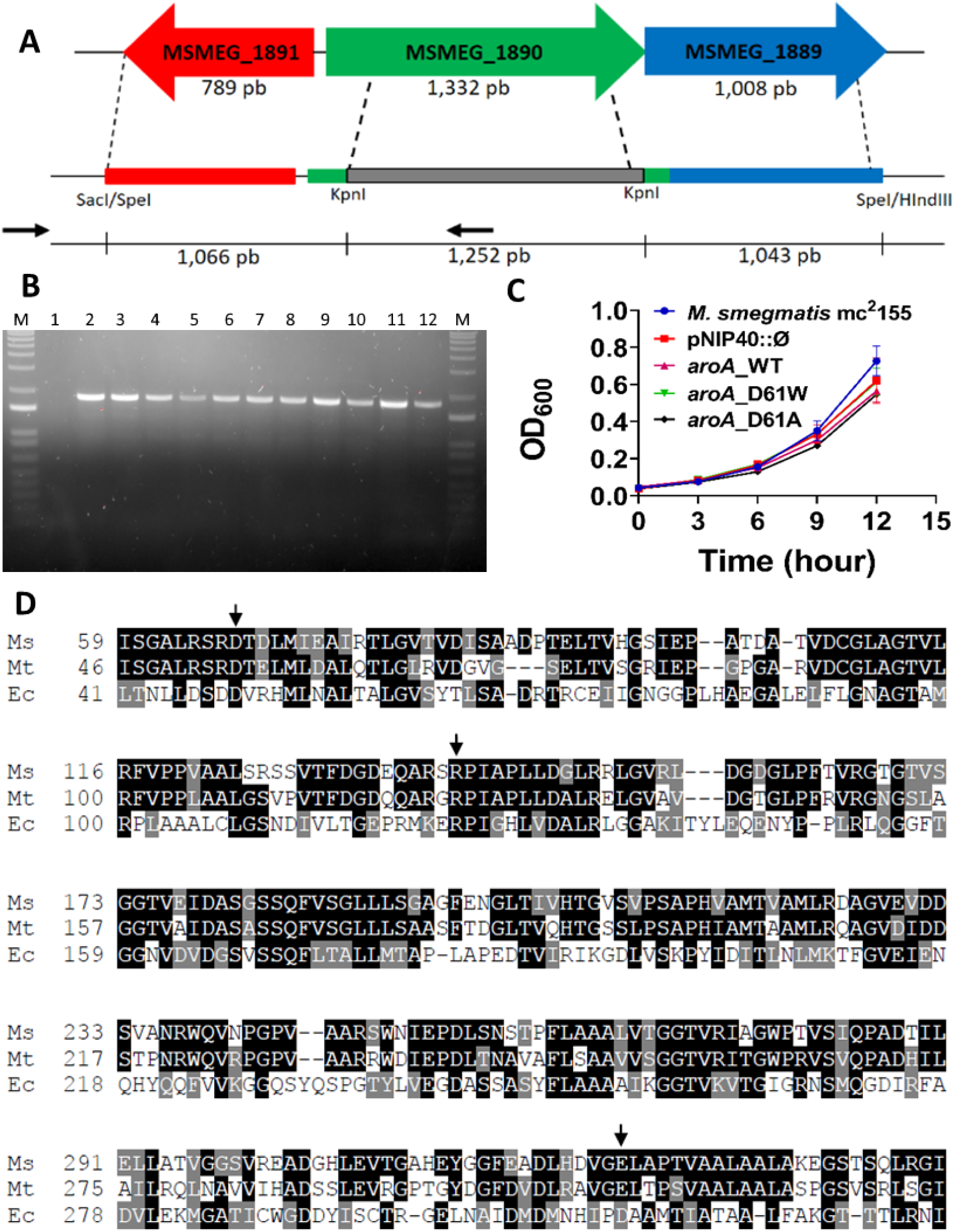
*aroA* gene from *M. smegmatis* is essential for mycobacterial survival *in vitro*. **(A)** Schematic representation of the allelic exchange event in the *aroA* locus. Two putative genes (MSMEG_1891 and MSMEG_1889) flank the *aroA* gene (MSMEG_1890) of *M. smegmatis*. The Allelic Exchange Sequences (AESs) were designed to maintain possible transcriptional and translation regulatory sequences of these two genes. The *aroA* gene was disrupted by the insertion of a kanamycin resistance cassette (1,252 bp), which was also used as a selective marker for homologous recombination. The position of primers used in PCR reactions described in (B) are indicated by black arrows. **(B)** PCR confirming the interruption of the *aroA* gene in merodiploid strains carrying the WT, D61A, or D61W mutant copies. Genomic DNA extracted from white colonies, and a pair of primers specific for the interrupted gene were used for this reaction. A band of 1,813 bp was expected for allelic exchange mutants. Lane M: 1kb plus DNA ladder (Invitrogen). Lane 1: *M. smegmatis* mc²155 genomic DNA (negative control). Lanes 2-8: strains carrying the WT copy of *aroA* gene. Lanes 9-12: strains carrying the D61A (9-10) or D61W (11-12) mutant copies of *aroA* gene. **(C)** Mutations in the Asp61 residue of *Ms*EPSPS does not impair mycobacterial survival and growth *in vitro*. Strains carrying mutations D61A and D61W in the *Ms*EPSPS enzyme were grown for 12 h in LB medium, under aerobic conditions, and aliquots were taken each 3 h for optical density measurement at 600 nm (OD_600_). Strains carrying the WT *aroA* gene or the empty complementation vector (pNIP::Ø), as well as the reference *M. smegmatis mc²155*, were used as growth references. Error bars are standard deviation (SD) of three biological replicates. **(D)** Sequence alignment of EPSPS enzymes from *E. coli* CVM N33429PS (Ec), *M. tuberculosis* H37Rv (Mt) and *M. smegmatis* mc^2^155 (Ms). Amino acid sequences were aligned by using T-Coffee and Boxshade. The enzyme from *M. smegmatis* shows 53% and 71% of identity with enzymes of *E. coli* and *M. tuberculosis*, respectively. Amino acids indicated by black arrows were chosen for mutagenesis.

### Aerobic growth curves of complemented strains

The complemented strains that were viable after the deletion of the wild-type *aroA* chromosomal copy were grown in LB medium + kanamycin + hygromycin, until reaching the early-log phase (OD_600_ ≈0.2). Then, cultures were diluted into fresh LB medium with antibiotics to a theoretical OD_600_ of 0.02, and divided (16 mL) in three conical tubes of 50 mL. Cultures were further incubated for 12 hours, at 37°C, under shaking (180 rpm) and aerobic conditions. Aliquots of 1 mL were taken every 3 hours and OD_600_ was measured. Results were expressed as mean ± standard deviation (SD) of three biological replicates.

### Gene knockdown by CRISPRi

Knockdown experiments using CRISPRi were performed on liquid and solid media. First, electrocompetent *M. smegmatis* cells were transformed with PLJR962 constructions containing the sgRNA scaffold coding sequences (see section “Construction of gene knockdown (KD) vectors”), and transformants were selected on solid LB with kanamycin. After three days of incubation, three isolated colonies were cultivated in 5 mL of LB for 48 h, at 37°C, under shaking (180 rpm). Next, cultures were diluted (1:200) in LB (100 mL) containing kanamycin, and further incubated to reach an OD_600_ of 0.2 (for growth curve) or 0.6 (for drop method on plates). For gene KD in liquid medium, cultures were further diluted in fresh medium (OD_600_ ≈0.02) containing kanamycin, equally divided (16 mL) in three conical tubes of 50 mL, with or without ATC, and grown for 24 h at 37°C. Samples (1 mL) were taken every 3 h. Results were expressed as mean ± standard deviation (SD) of three biological replicates. For gene KD in solid medium, drops of 5 µL were plated on solid LB containing kanamycin, with or without ATC. The first spot contained approximately 5,000 cells, and the other three subsequent spots were tenfold serially diluted. Plates were incubated for 3-4 days at 37°C. A negative (PLJR962::Ø) and a positive control (PLJR962::*mmpL3*) were employed for each condition. Additionally, KD experiments were performed in 7H9/7H10 media in the presence, or absence of aromatic amino acids.

### Protein Extraction of *M. smegmatis*

For each sample of gene KD in liquid medium, total protein was extracted in 0h and 18h, as previously described [25, 26]. Cellular pellets were washed twice with 10 mM Tris-HCl pH 8.0 and then collected by centrifugation (4,000 rpm, 15 min, 4°C) (Hitachi himac CR21G centrifuge) and resuspended in 2 mL of the same buffer. Cells were disrupted by sonication (10 pulses of 10 s, with intervals of 1 min on ice at 21% of amplitude) using the Sonics Vibra Cell equipment (High Intensity Ultrasonic Processor, 750 Watt model) with a 13 mm probe, centrifuged (13,000 rpm, 30 min, 4°C) and the supernatant (soluble proteins) was stored at −80°C.

### Western Blots

Anti-*M. tuberculosis* EPSPS (*Mt*EPSPS) mouse polyclonal antibody was produced immunizing a mouse with 50 mg of purified recombinant *Mt*EPSPS containing Freund’s incomplete adjuvant (Sigma-Aldrich, USA) (total volume of 100 µL) by subcutaneous route, followed by a booster injection after one month. The mouse was euthanized by deep isoflurane inhalation one month later, and blood was collected by the descendant aorta. Serum was separated by centrifugation at 10,000 x g for 10 min, aliquoted, and stored at −80°C [26]. The western blot was performed in triplicate. Approximately 30 µg of *M. smegmatis* proteins from detergent fraction were boiled at 70°C for 10 min, loaded on 12% sodium dodecyl sulfate polyacrylamide gels (SDS-PAGE), and transferred to nitrocellulose membranes (Merck Millipore, Ltd-Ireland) in Buffer Tris 25 mM, glycine 192 mM pH 8.8 and methanol 20% for 4h at 70 v. After transfer, the membrane was blocked with 5% non-fat dried milk (Santa Cruz Biotechnology, USA), 0.05% tween-20 (Sigma-Aldrich, USA) in TBS (T-TBS) (2h, 4°C) and probed with anti-*Mt*EPSPS polyclonal mouse antibody in a 1:500 dilution (overnight at 4°C). Membranes were washed three times with T-TBS, and alkaline phosphatase-conjugated anti-mouse secondary antibody (Invitrogen, USA) was used at a dilution of 1:5,000 [27]. Chemiluminescent substrate (Novex by Life Technologies, USA) was used for detection with ChemiDoc (Bio-Rad, USA).

### Overexpression of WT and mutants of *M. smegmatis*

*E. coli* cells were transformed with recombinant pET-23a(+) plasmids carrying the WT or mutants (R134A, E321N or D61W) of *aroA* gene, and selected on solid LB with ampicillin. A single colony was grown in LB medium (5 mL) with antibiotic, at 37°C, O/N. Pre-cultivated inocula were then diluted (1:1000) in fresh LB (for WT, R134A and E321N) or Terrific Broth (TB) media (for D61W), containing ampicillin. After reaching an OD_600_ of 0.4-0.6, cultures were grown for 23 h at 37°C, under shaking (180 rpm) and aerobic conditions. Protein expression was achieved without isopropyl β-D-1-thiogalactoside (IPTG) induction. Cells were harvested by centrifugation (11,800 x g for 30 min, at 4°C), and stored at - 20°C. As a negative control of the expression, the same procedure was employed for *E. coli* cells carrying pET-23a(+) without the *aroA* gene (pET23a(+)::Ø). The expression of soluble proteins was confirmed by 12% SDS-PAGE stained with Coomassie Brilliant Blue.

### Purification of recombinant proteins by liquid chromatography

Recombinant WT and mutant proteins were purified using two or three chromatographic steps. All purification steps were carried out in an ÄKTA system (GE Healthcare® Life Sciences) at 4°C. Approximately 3.2 g of cells overproducing each protein were collected. Cells were suspended in 25 mL of 50 mM Tris–HCl pH 7.8 (buffer A), and incubated for 30 min in the presence of 0.2 mg/mL lysozyme (Sigma-Aldrich), under slow stirring. Cells were disrupted by sonication (4 pulses of 20 s, with intervals of 1 min on ice, at 60% of amplitude). Cell debris were removed by centrifugation (11,800 x g for 60 min, at 4 °C). The supernatant was incubated with 1% (w/v) of streptomycin sulphate (Sigma-Aldrich) for 30 min at 4°C, under gently stirring, and centrifuged. The supernatant was dialyzed twice against 2 L of buffer A, using a dialysis tubing with a cutoff filter of 12–14 kDa. The samples were clarified by centrifugation and loaded onto a Q-Sepharose Fast Flow (GE Healthcare^®^ Life Sciences) column, pre-equilibrated with buffer A. Adsorbed proteins were eluted by a 20 column volume (CV) linear gradient (0 - 100%) of 50 mM Tris–HCl NaCl 1M pH 7.8 (buffer B), at 1 mL/min flow rate. Protein elution was monitored by UV detection at 215, 254, and 280 nm. Eluted fractions containing the protein of interest were pooled and ammonium sulphate was added to a final concentration of 1 M. After an incubation period of 30 min at 4°C, and subsequent centrifugation, the supernatant was loaded on a HiLoad 16/10 Phenyl Sepharose HP (GE Healthcare^®^ Life Sciences) column, pre-equilibrated with 50 mM Tris-HCl (NH_4_)_2_SO_4_ 1 M pH 7.8 (buffer C). Proteins were eluted by a 20 CV linear gradient (100 - 0% ammonium sulphate) in buffer A, at 1 mL/min flow rate. For WT and R134A mutant, a Mono Q HR 16/10 (GE Healthcare® Life Sciences) column was used as a third step. Protein fractions eluted from the second column were pooled, centrifuged and loaded into the last column, pre-equilibrated with buffer A. Proteins were eluted by a 15 CV linear gradient in buffer B (0 - 100%), at 2 mL/min flow rate, pooled and dialyzed against buffer A, and finally stored at - 80°C. All protein fractions were analyzed by 12% SDS-PAGE stained with Coomassie Brilliant Blue. Protein homogeneity above 95% was checked by densitometry in a GelDoc (Bio-Rad) equipment. Protein concentration was determined by BCA method (Thermo Scientific Pierce™ BCA protein Assay Kit).

### Protein identification by LC-MS/MS

Recombinant *Ms*EPSPS enzymes were precipitated with chloroform/methanol [28]. Pellets were resuspended in 100 mM Tris-HCl pH 7.0 containing 8 M urea (Affymetrix USB) and disulfide bonds were reduced in 5 mM dithiothreitol (DTT) (Ludwig Biotec) for 20 min at 37°C. After that, cysteine residues were alkylated with 25 mM iodoacetamide (IAM) (Sigma-Aldrich) for 20 min at room temperature in the dark. Urea was diluted to 2 M with 100 mM Tris-HCl pH 7.0 and trypsin (Promega) was added at a mass ratio of 1:100 (trypsin:protein). Digestion was incubated overnight at 37°C. Formic acid (Merck) was added to end the reaction (5% v/v, final concentration). Tryptic peptides were then separated in a reversed phase C18 (5 µm ODS-AQ C18, Yamamura Chemical Lab) column using a nanoUPLC (nanoLC Ultra 1D plus, Eksigent, USA) and eluted (400 nL/min) with acetonitrile gradient (5%-80%) (LiChrosolv^®^, Merck) with 0.1% formic acid. Eluting peptide fragments were ionized by electrospray ionization and analyzed on a LTQ-XL Orbitrap Discovery hybrid instrument (Thermo Fisher Scientific). The LC–MS/MS procedure was performed according to the data-dependent acquisition (DDA) method. Precursors were collected from 400-1600 m/z at 30,000 resolution in the Orbitrap and the eight most abundant ions per scan were selected to collision-induced dissociation (CID), using helium as the collision gas, in the ion trap. Raw files were searched in the PatternLab for Proteomics platform [29] with a database containing forward and reverse *E. coli* BL21-DE3 reference proteome and *Ms*EPSPS WT and mutants sequences using Comet [30]. Carbamidomethyl was set as a fixed modification. Search results were filtered to a false discovery rate of 1% through the module Search Engine Processor from PatternLab for Proteomics.

### EPSPS enzyme activity assays

Recombinant *Ms*EPSPS enzymes were assayed in the forward direction, using a continuous spectrophotometric rate assay [31, 32]. Enzyme activity was measured in a coupled assay with purine nucleoside phosphorylase from *M. tuberculosis* (*Mt*PNP; EC 2.4.2.1), and 2-amino-6-mercapto-7-methylpurine ribonucleoside (MESG), which was synthesized according to a published protocol [33] (Supplemental Material - Fig. S1). All activity assays were performed in 100 mM Tris–HCl buffer, pH 7.8, at 25°C for 3 min, using 138 nM of *Mt*PNP and 1.7 nM of *Ms*EPSPS. Apparent steady-state kinetic constants were determined by monitoring the WT and mutant EPSPS activities at varying concentrations of 2-phosphoenolpyruvate (PEP – Sigma-Aldrich), and fixed-saturating concentrations of shikimate-3-phosphate (S3P - Sigma-Aldrich) (Supplemental Material – Table S1). All measurements were performed in a 1.0 cm path length quartz cuvette, in duplicate, and the rate of inorganic phosphate (Pi) production was measured in a UV/Vis spectrophotometer (Shimadzu). Steady-state kinetic constants were obtained by non-linear regression analysis of the kinetic data fitted to the Michaelis-Menten equation (*v* = *V*_max_ x [S]/(*K_m_* + [S])) using the SigmaPlot 14.0 software (SPSS, Inc). The *k*_cat_ values were calculated using the following *k*_cat_ equation (*k*_cat_ = *V*_max_/[E]t).

## RESULTS

### *In vitro* essentiality of *aroA* gene from *M. smegmatis*

We knocked out the *aroA* gene from *M. smegmatis* to evaluate its essentiality *in vitro.* First, a set of merodiploid strains, holding an extra copy of the WT or mutants *aroA* genes (D61A, D61W, E321N, and R134A), received a plasmid carrying the allelic exchange substrate (pPR27::KO_*aroA*), which was confirmed by the presence of bright yellow colonies after catechol addition (see Methods, section “Knockout of the aroA gene from *M. smegmatis* and gene complementation”). Only three independent yellow colonies were grown under permissive conditions, and then submitted to counterselective pressures (growth temperature of 39°C and 2% sucrose) on solid LB medium. From an inoculum of ≈10^7^ CFUs per plate, several white colonies (Kan^R^, Hyg^R^, XylE^-^, Suc^R^), holding the WT *aroA* extra copy, were obtained. Only three colonies were observed from the strain carrying no extra copy of *aroA* (pNIP::Ø), but they revealed to be yellow after catechol testing (Kan^R^, Hyg^R^, XylE^+^, Suc^R^). This result suggests that in the absence of a functional copy of the *aroA* gene the mycobacteria is unable to survive, confirming the essentiality of this gene *in vitro*. The same result was observed in 7H10 medium.

We conducted similar experiments with merodiploid strains containing an extra copy of *aroA* gene (WT, D61A, D61W, R134A and E321N). In both LB and 7H10 media, only the WT strain and strains carrying mutations in the aspartic acid residue (D61A and D61W) survived the allelic exchange event. To confirm the DCO event, a PCR reaction was carried out using genomic DNA extracted from each of the white colonies obtained (Fig. 2B).

### Mutations in the Asp61 residue of *Ms*EPSPS enzyme enables mycobacterial growth

Growth curves were performed to evaluate the impact of mutations in the aspartic acid 61 residue (D61A or D61W) of *Ms*EPSPS on bacilli grown in LB medium. We found no differences in the growth of D61 mutants, when compared to control strains (Fig. 2C). This suggests that the replacement of this hydrophilic amino acid by the hydrophobic alanine or tryptophan residues does not abolish the *Ms*EPSPS activity inside cells.

### Expression, purification and identification of recombinant *Ms*EPSPSs

The overexpression of recombinant *Ms*EPSPS proteins (WT, D61W, R134A and E321N) in the soluble fraction was confirmed by SDS-PAGE, with an apparent molecular mass of 46 kDa. Homogeneous preparations were obtained using a 3-step protocol for both *Ms*EPSPS WT and D61W, whereas a 2-step protocol was employed for mutants R134A and E321N (Table 2, Supplemental Material – Fig. S2). Recombinant *Ms*EPSPS WT, D61W, E321N and R134A mutants were submitted to trypsin digestion and peptides were analyzed by LC-MS/MS. A coverage of approximately 90% was obtained for each protein with 85, 97, 80 and 74 unique peptides identified, respectively. Furthermore, it was possible to identify and validate all point mutations (Supplemental Material - Fig. S3-5).

**Table 2.** Purification yield of recombinant *Ms*EPSPS enzymes.

| *MsEPSPS*<br>Enzymes<sup>a</sup> | Column<sup>b</sup> | Protein<br>concentration<br>(mg/mL) | Eluted<br>volume<br>(mL) | Total<br>protein<br>(mg) | Yield<br>(%) | Homogeneity<br>(%) |
| --- | --- | --- | --- | --- | --- | --- |
| WT | First | 13.4 | 25 | 334.4 | 20.7 | 97.3 |
|  | Last | 3.6 | 19 | 69.4 |  |  |
| D61W | First | 6.4 | 25 | 160.7 | 8.5 | 96.8 |
|  | Last | 0.7 | 20 | 13.7 |  |  |
| R134A | First | 20.4 | 25 | 510.7 | 11.2 | 98.4 |
|  | Last | 3.0 | 19 | 57.1 |  |  |
| E321N | First | 13.2 | 25 | 330.0 | 17.3 | 100 |
|  | Last | 0.6 | 96 | 57.2 |  |  |

<sup>a</sup>Recombinant wild-type (WT) or mutants EPSPS enzymes from *M. smegmatis*.
<sup>b</sup>Chromatographic column used in the first or last step of the purification protocol.

### Kinetic parameters of WT and mutant EPSPS enzymes

EPSPS enzymes are known to catalyze the transfer of the carboxyvinyl portion of the phosphoenolpyruvate substrate (PEP) to the carbon-5 hydroxyl group of shikimate-3-phosphate (S3P), forming the enolpyruvylshikimate-3-phosphate (EPSP) product and releasing inorganic phosphate (Pi). To determine the kinetic parameters of WT and mutant enzymes, we performed a coupled assay using *Mt*PNP and MESG. The dependence of initial velocity on PEP as a variable substrate at fixed-saturating S3P concentration (see Supplemental Material – Table S1) followed hyperbolic Michaelis-Menten kinetics. The apparent steady-state kinetic parameters for WT and mutant *Ms*EPSPS enzymes are presented in Table 3.

**Table 3.** Apparent steady-state kinetic parameters for *Ms*EPSPS enzymes.

| *MsEPSPS*<br>Enzymes<sup>a</sup> | $K_m$ (μM) | $k_{cat}$ (s<sup>-1</sup>) | $k_{cat}/K_m$ (M<sup>-1</sup>s<sup>-1</sup>) |
| --- | --- | --- | --- |
| **WT** | 88 ± 11 | 0.5530 ± 0.0185 | 6.28 E+03 ± 813 |
| **D61W** | 1014 ± 975 | 0.1075 ± 0.0608 | 1.06 E+02 ± 118 |
| **R134A** | 3676 ± 1007 | 0.1843 ± 0.0330 | 5.01 E+01 ± 16 |
| **E321N** | 3081 ± 808 | 0.4378 ± 0.0469 | 1.42 E+02 ± 40 |

<sup>a</sup>Recombinant EPSPS enzymes from *M. smegmatis*: WT and point mutants. S3P was used at saturating concentrations (Supplemental Material – Table S1) and PEP as a variable substrate in the enzymatic assay. All reactions were performed in duplicate.

### *aroA* silencing with CRISPRi in *M. smegmatis*

The vulnerability of *aroA* from *M. smegmatis* was assessed by using CRISPRi in different growth contexts. Using an in-house script written in Python, twelve targets were found in the non-template strand (NT) of *aroA* coding sequence. Three distinct sequences next to functional PAMs (5’-NAGCAT-3’, 5’-NNAGGAT-3’, and 5’-NNAGCAG-3’) and located at the first half of the gene (Fig. 1) were chosen to be targeted by three sgRNAs (named PAM1, PAM2, and PAM3). The vulnerability of this gene was evaluated in both rich media (solid and liquid LB – Fig. 3A-D) and defined media (solid 7H10 and liquid 7H9 – Fig. 3E-H) in the presence or absence of ATC 100 ng/mL, using the vulnerable *mmpL3* gene as positive control. We did not observe any difference in growth in the presence or absence of ATC in both solid and liquid rich media (Fig. 3B-D). In contrast, with all target sequences tested (adjacent to PAM 1 to 3) in solid and liquid defined media, we observed a decrease in bacterial growth from 15h in the presence of ATC, indicating that *aroA* gene silencing leads to a bacterial growth perturbation in poor nutrients media (Fig. 3F-H).

**Figure 3.**
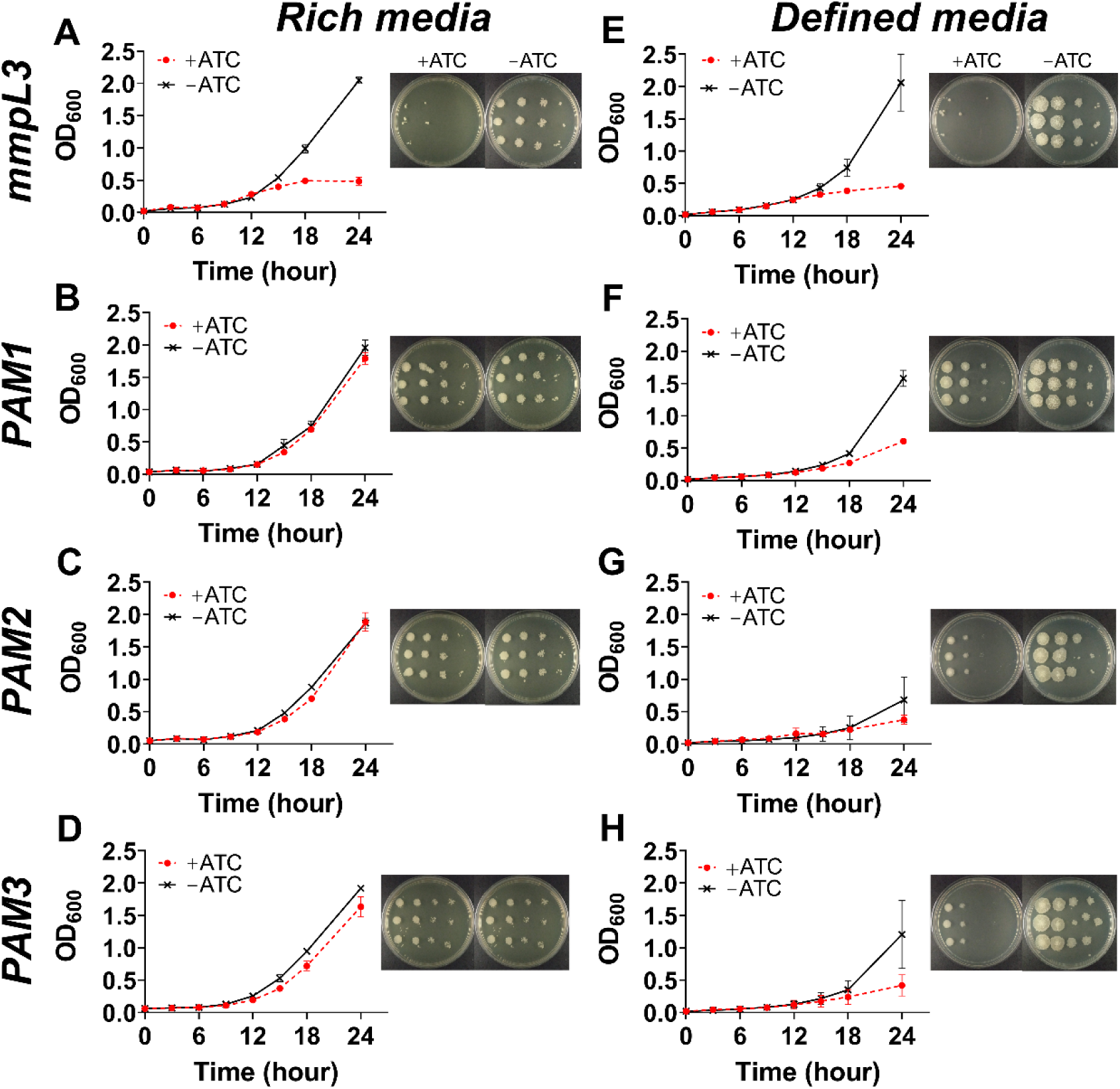
Knockdown of *aroA* gene from *M. smegmatis* produces a growth perturbation *in vitro.* **(A-D)** Growth in rich media (solid and liquid LB). **(E-H)** Growth in defined media (solid 7H10 and liquid 7H9). *M. smegmatis* growth curves and dilution spots in the presence or absence of anhydrotetracycline (ATC) (100 ng/mL) for the control gene *mmpL3* **(A and E)** and the *aroA* gene at three different locations adjacent to PAM1, PAM2 and PAM3 **(B-D and F-H)**.

Next, we supplemented solid defined medium (7H10) with aromatic amino acids (L-tryptophan + L-phenylalanine + L-tyrosine), which are end products of the Shikimate pathway (Fig. 4A), and repeated the *aroA* knockdown using the CRISPRi system. Interestingly, we did not observe any difference in growth in the presence or in the absence of ATC (Fig. 4B-D).

**Figure 4.**
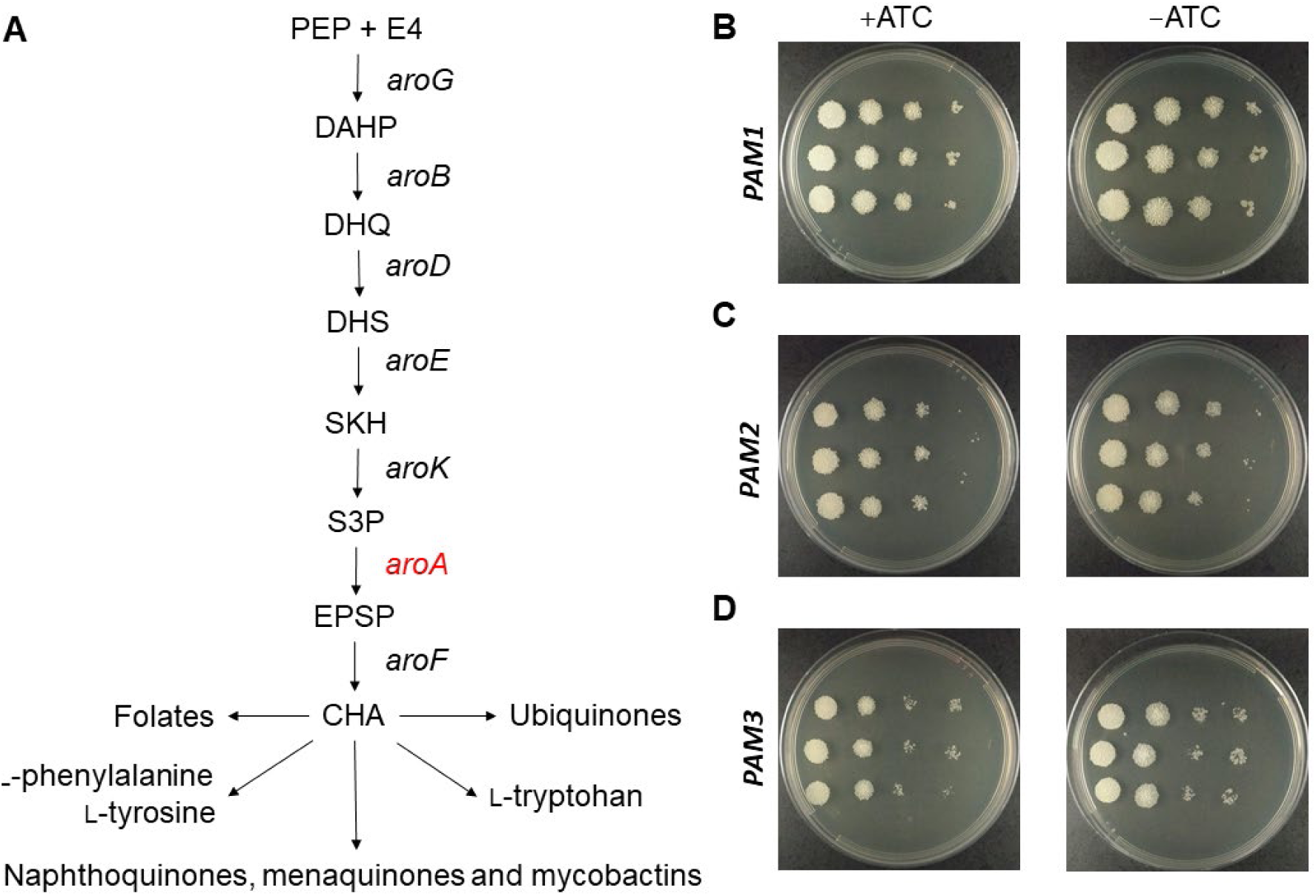
Rescued *M. smegmatis* strains in the presence of the aromatic amino acids. **(A)** Schematic representation of shikimate pathway and its end products. **(B-D)** *M. smegmatis* growth in the presence or absence of ATC (100 ng/mL) in solid defined medium (7H10) supplemented with aromatic amino acids (L-phenylalanine, L-tyrosine and L-tryptophan).

## DISCUSSION

Target validation is a required part of any effort to develop new chemotherapeutic agents based on rational-drug design. Essentiality for growth and/or survival is a critical feature of a target, as the chemical inhibition of non-essential gene products are not expected to kill the infective agent and hence to achieve the desired therapeutic outcome. Here, by performing an allelic exchange-based gene knockout experiment, we show that the *aroA*-encoded EPSPS gene product is essential for *M. smegmatis* growth *in vitro* (Fig. 2). This agrees with previous reports based on transposon-mediated mutagenesis that the orthologous gene from *M. tuberculosis* is also essential [34].

Previously, the *aroK* gene from *M. tuberculosis*, which encodes shikimate kinase (*Mt*SK), was also found to be essential [13]. Interestingly, the supplementation of neither the end product of the pathway, chorismate, nor aromatic amino acids (tyrosine, tryptophan and phenylalanine) was sufficient to allow growth of *aroK* mutants. It was suggested that *Mt*SK, like *aroK*-encoded SK from *Escherichia coli* [35], could have other functions unrelated to the shikimate kinase activity [13]. We thus evaluated whether the EPSPS activity of *aroA* protein product is responsible for *aroA* gene essentiality. To do so, we constructed four merodiploid strains containing extra copies of *aroA* encoding point mutants of EPSPS (D61A, D61W, R134A and E321N).

The selection of these residues was based on a previous experimental work on EPSPS from *E. coli* (*Ec*EPSPS) [36] and on computational studies of EPSPS from *M. tuberculosis* (*Mt*EPSPS) [37]. It was already demonstrated that a mutation in the aspartic acid-49 (D49) residue to an alanine leads to a reduction of 24,000 times in the specific activity of the enzyme from *E. coli*. The reasons for that are still unclear, but the authors hypothesized an indirect effect on the lysine-22 (K22) residue, which is known to participate directly in catalysis. On the other hand, the residues arginine-124 (R124) and aspartic acid-313 (D313), which are near to the PEP binding site, are directly involved in the catalytic reaction. When those residues were mutated to an alanine and glutamic acid, respectively, the enzymes had their catalytic activity reduced to around 5,000 and 20,000 times, showing that residues R124 and D313 are critical to the correct function of the *Ec*EPSPS enzyme [36]. In addition, *in silico* predictions using the enzyme of *M. tuberculosis* suggested that changing the aspartic acid-54 (D54) residue (which corresponds to D49 in *Ec*EPSPS and D61 in *Ms*EPSPS enzymes) to an alanine (D54A) or tryptophan (D54W) should cause a significant impact on the protein stability and, consequently, a negative impact on the enzyme’s activity [37].

To confirm the expected impact of these point mutations on *Ms*EPSPS activity, we cloned, expressed and purified mutants D61W, R134A and E321N. The kinetic properties of the wild-type recombinant enzyme (WT) and three mutants were measured and compared (Table 3). The *K*_m_ for the substrate PEP in mutant forms of *Ms*EPSPS increased from 11.5 up to 42 times when compared to WT enzyme. These results suggest an increased overall dissociation constant for PEP substrate binding to mutant proteins at fixed-saturating concentrations of S3P. The impact on enzyme turnover (*k_cat_*) ranged from 1.2-fold (E321N) to 5.1-fold (D61W) decrease (Table 3). Accordingly, more pronounced effects on the catalytic efficiencies (*k*_cat_/*K*_m_) of mutants were observed. We found a reduction of 44- (E321N), 59- (D61W) and 125-fold (R134A) in *k*_cat_/*K*_m_ for these enzymes. These reductions in the apparent second-order rate constants suggest lower association rate constants for PEP substrate binding to *Ms*EPSPS enzyme. Therefore, we can conclude that mutations in these specific residues affect directly the catalytic efficiency of the *Ms*EPSPS, although to a lesser extent than expected, based on previous studies with orthologs [36].

In the intracellular environment, however, the impact of each mutation in the cell metabolism was different. The replacement of aspartate-61 with hydrophobic residues (alanine or tryptophan) was not lethal (Fig. 2B) and did not impart in any growth defect of *M. smegmatis* (Fig. 2C). This may suggest that inside cells aspartate-61 is dispensable for *Ms*EPSPS activity, or the 59-fold decrease in the *k*_cat_/*K*_m_ value for the PEP substrate is not sufficient to impair cell growth. On the other hand, we were not able to retrieve viable colonies after knocking out the WT *aroA* gene from merodiploid strains containing an extra copy of *aroA* gene encoding *Ms*EPSPS R134A or E321N mutants. These results suggest that both mutations are lethal under the conditions tested and corroborate the proposition that *aroA* essentiality is causally linked to EPSPS activity. As pointed out by some of us, establishing a causal link between gene essentiality and the biological function of its protein product under scrutiny should be an indispensable step in target validation for drug development [24]. This view is reinforced by the growing number of proteins found to exhibit multiple and unrelated tasks, the so-called moonlighting proteins [38].

Next, we addressed the issue of target vulnerability. A target should not only be essential but also vulnerable, otherwise chances are low to develop bioactive compounds that effectively kill or impart growth defects on infective agents. To evaluate *Ms*EPSPS vulnerability, we performed gene knockdown experiments using the CRISPRi system developed for mycobacteria [18]. Using an in-house Python script, we selected target sequences adjacent to PAM motifs whose repression strengths were characterized previously [18]. The experiments were conducted in both rich (LB, Fig. 3A-D) and defined (7H9 or 7H10, Fig. 3E-H) media, with markedly different results. As expected, the sgRNA control targeting the *mmpL3* gene was found to be vulnerable in both nutritional conditions, either in liquid or solid media. Silencing *mmpL3* caused a cessation of bacterial growth by 15 h, and using the drop method on plates, it was observed a reduction of at least 1000-fold in the CFU counting (Fig. 3A and 3E). This gene codes for the mycobacterial membrane MmpL3 protein, which is responsible for trehalose monomycolate transportation through the cell inner membrane [39]. In *M. tuberculosis* and *M. smegmatis*, it was shown that silencing *mmpL3* expression disrupts bacterial growth [18, 40], leading to accumulation of TDM and cell death [41]. Differently from MmpL3, *Ms*EPSPS was found to be vulnerable only in defined media and in the presence of ATC, irrespective of PAM’s repression strength. Silencing *aroA* gene caused an impairment of the bacterial growth after 18 h, but not a complete cessation (Fig. 3F-H). This suggests that the suppression of *Ms*EPSPS expression observed (Supplemental Material – Fig. S6-7) does not cause bacterial killing, but rather, growth impairment (Fig. 3F-H). Moreover, supplementation with aromatic amino acids (L-phenylalanine, L-tyrosine and L-tryptophan) is sufficient to rescue the growth impairment of *aroA*-silenced strain (Fig. 4B-D).

Drawing a parallel between our results of *aroA* gene knockout and *aroA* gene knockdown with CRISPRi, it is evident that both experimental approaches serve to understand different biological phenomena. By performing gene knockout and complementation experiments, we found that *aroA* gene of *M. smegmatis* is essential regardless of the nutritional context (Fig. 2A-C, Fig. S8). This is an evidence that this bacterium is unable to grow in the complete absence of *Ms*EPSPS, most likely because of its incapacity of producing chorismate [13]. This metabolite is a precursor for the synthesis of folates, aromatic amino acids, ubiquinones, naphtoquinones, menaquinones and mycobactins (Fig. 4A), being indispensable for bacterial metabolism. On the other hand, even after resulting in markedly reduced levels of *Ms*EPSPS (Fig. S6 and S7), *aroA* gene knockdown using the CRISPRi system impaired bacterial growth only in a poor nutritional context (Fig. 3), and supplementation of aromatic amino acids was sufficient to restore normal growth (Fig. 4B-D). Presumably, very low levels of *Ms*EPSPS can support the operation of the shikimate pathway to the level required to produce most of the metabolic end products that have chorismate as a precursor compound, except for the aromatic amino acids. The implications of our results are twofold. By growing *aroA*-knocked down bacteria in different nutritional conditions, rich medium, defined medium and defined medium with supplementation, we were able to characterize *Ms*EPSPS vulnerability in more detail. In a context of low availability of aromatic amino acids, *Ms*EPSPS can be considered a vulnerable target. Otherwise, under the presence of appropriate levels of L-phenylalanine, L-tyrosine and L-tryptophan, our experiments suggest *Ms*EPSPS would not be a vulnerable target, as the incomplete inhibition of its activity by antimicrobial agents is not expected to abrogate the synthesis of the other chorismate-dependent end products and lead to growth impairment and cell death. Moreover, the results we obtained using both gene knockout and gene knockdown approaches raises a cautionary note to the use of gene knockdown experiments to infer gene essentiality. As was the case with *Ms*EPSPS, the presence of very low protein levels, undetectable by means of Western Blot assays (Figs. S6 and S7), can lead to a completely different phenotype when compared to that obtained under the complete absence of the same protein, as in knocked out strains (Figs. 2A-C and S8), and consequently to a misappreciation of gene essentiality.

## Acknowledgments

We thank Sara Fortune and Jeremy Rock for providing the pLJR962 plasmid and Hector Morbidoni and Luis Saraiva Timmers for insightful discussions. C.V.B., P.M. and L.A.B. would like to acknowledge financial support given by CNPq/FAPERGS/CAPES/BNDES to the National Institute of Science and Technology on Tuberculosis (INCT-TB), Brazil (grant numbers: 421703-2017-2/17-1265-8/14.2.0914.1). C.V.B. (310344/2016-6), P.M. (305203/2018-5) and L.A.B. (520182/99-5) are research career awardees of the National Council for Scientific and Technological Development of Brazil (CNPq). This study was financed in part by the Coordenação de Aperfeiçoamento de Pessoal de Nível Superior—Brasil (CAPES)—Finance Code 001.

## Supplemental Material

**Figure S1.**
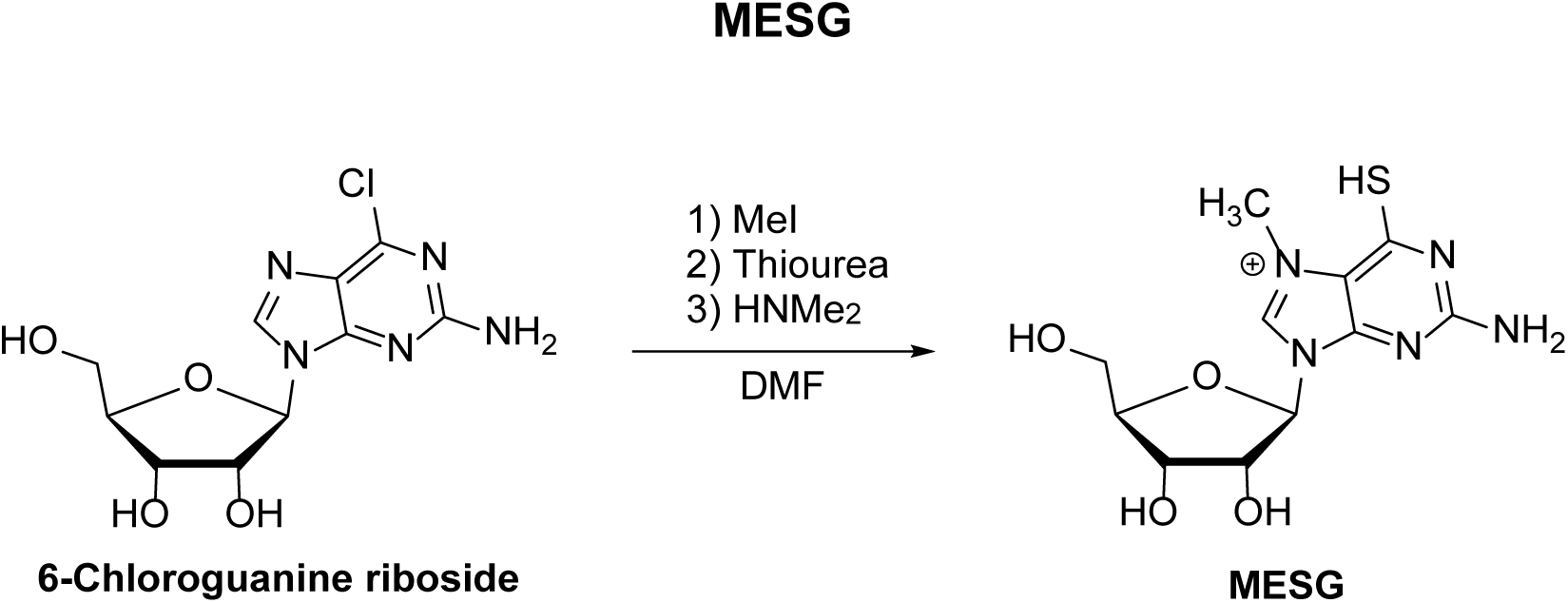
**MESG synthesis**. In a two-neck round bottom flask, under argon atmosphere, 6-chloro-guanine riboside (4.00 g, 13.25 mmol) was dissolved in dry dimethylformamide (10 mL). Then, methyl iodide (4 mL, 64.25 mmol) was added and the mixture was stirred overnight at 30 °C. Excess methyl iodide was removed under vacuum together with part of DMF. Then, to the residual mixture thiourea (2.00 g, 26.27 mmol) was added under an argon atmosphere and the mixture was stirred for an additional hour. Afterwards, the solution was neutralized with pure dimethylamine slowly added dropwise. The mixture was directly poured into stirred acetone (500 mL) to give a yellow precipitate which was further chromatographed in silica eluted with ethyl acetate/1-propanol/water (5:2:1; v/v) yielding 0,620g (30%) of **MESG**. The compound was dried to a yellow solid and stored desiccated at −80 °C. ^13^C NMR (D_2_O) δ (ppm): 174.2, 156.0, 146.6, 141.1, 119.6, 90.8, 86.2, 75.1, 70.1, 61.3, 35.4. HRMS (ESI): calc. for [C_11_H_16_N_5_O_4_S+H]^+^: 314.0918; obt: 314.0916.^1^

## Supplementary Results

### MESG synthesis

High-resolution mass spectra (HRMS) were obtained on an LTQ Orbitrap Discovery mass spectrometer (Thermo Fisher Scientific). This system combines an LTQ XL linear ion-trap mass spectrometer and an Orbitrap mass analyzer. The analyses were performed through the direct infusion of the sample in MeOH/H_2_O (1:1) with 0.1% formic acid (flow rate 10 mL/min) in a positive-ion mode using electrospray ionization (ESI). For elemental composition, calculations used the specific tool included in the Qual Browser module of Xcalibur (Thermo Fisher Scientific, release 2.0.7) software. ^13^C NMR spectra were acquired on an Avance III HD Bruker spectrometer (Pontifical Catholic University of Rio Grande do Sul); chemical shifts (δ) were expressed in parts per million (ppm) relative to TMS used as an internal standard.

**Table S1.** Fixed and varying concentrations of substrates used in kinetic assays of *Ms*EPSPS.

| EPSPS | Substrate | Fixed-saturating (μM) | Substrate | Varying range (μM) |
| --- | --- | --- | --- | --- |
| WT | S3P | 800 | PEP | 25 - 900 |
| D61W | S3P | 600 | PEP | 25 - 1200 |
| E321N | S3P | 600 | PEP | 25 - 10000 |
| R134A | S3P | 600 | PEP | 300 - 2900 |

**Figure S2.**
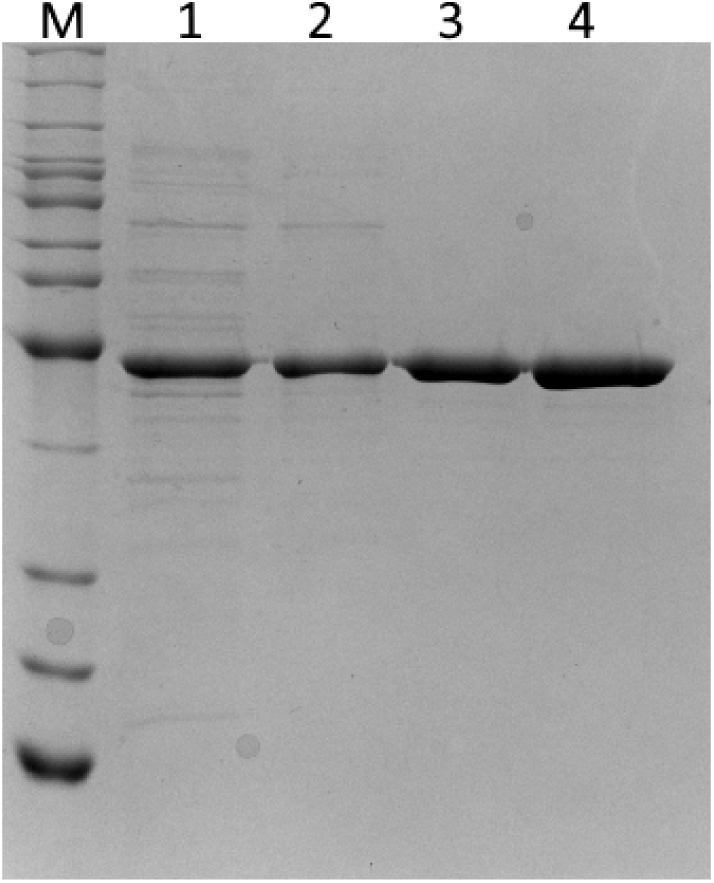
Representative SDS-PAGE from the purification steps of *Ms*EPSPS. Lane M: BenchMark Protein Leadder (Invitrogen). Lane 1: crude extract from soluble fraction of cell disruption. Lane 2: soluble fraction from the first column (Q-Sepharose Fast Flow). Lane 3: soluble fraction from the second column (Phenyl Sepharose HP) and Lane 4: soluble fraction from the third column (Mono Q HR 16/10).

**Figure S3.**
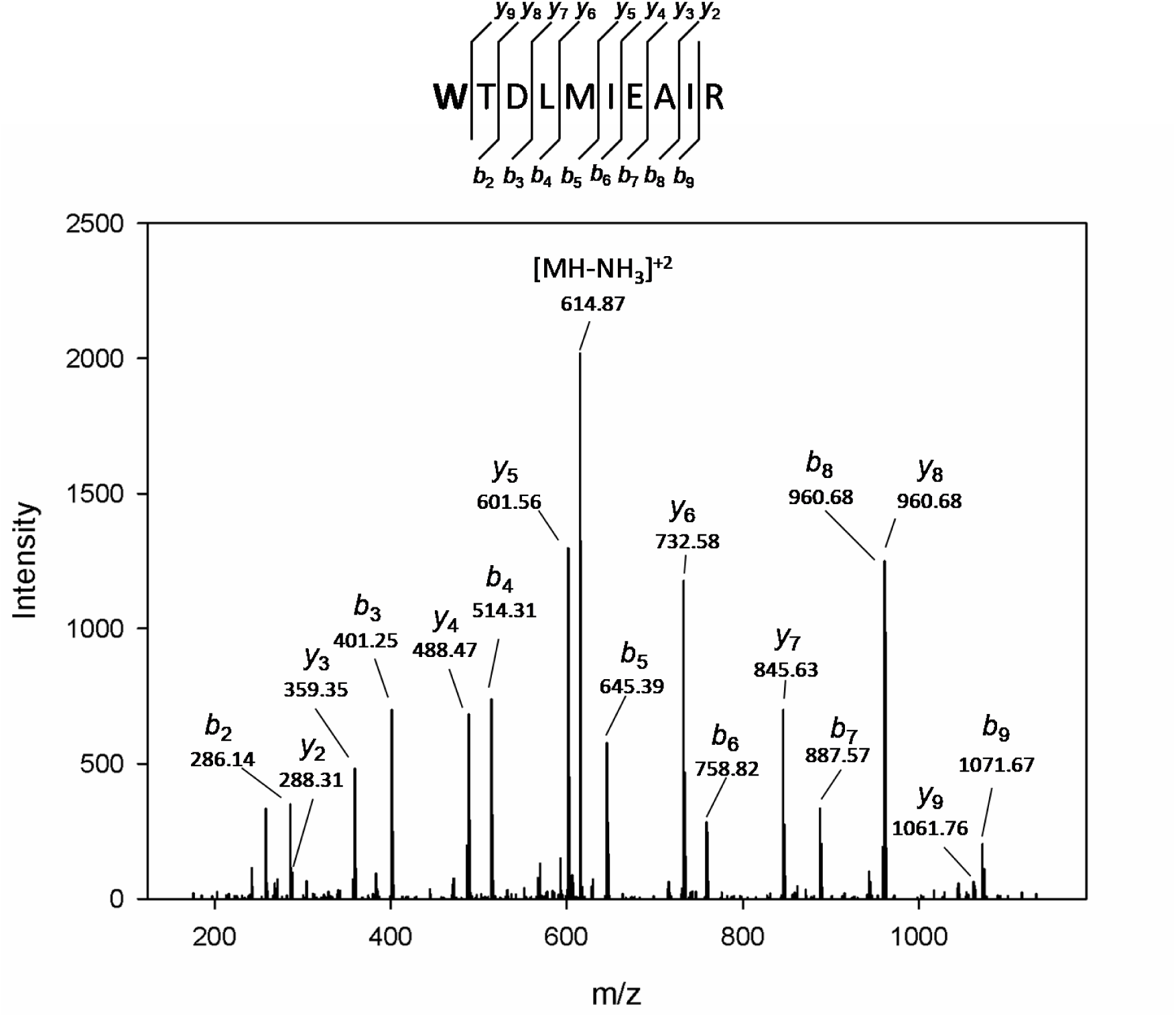
Representative spectra of peptide containing the D61W mutation obtained by LC-MS/MS of *Ms*EPSPS D61W protein. Peptide sequence: **W**TDLMIEAIR. Point mutation marked in **bold**. Fragment b- and y-ions and their neutral loses are indicated.

**Figure S4.**
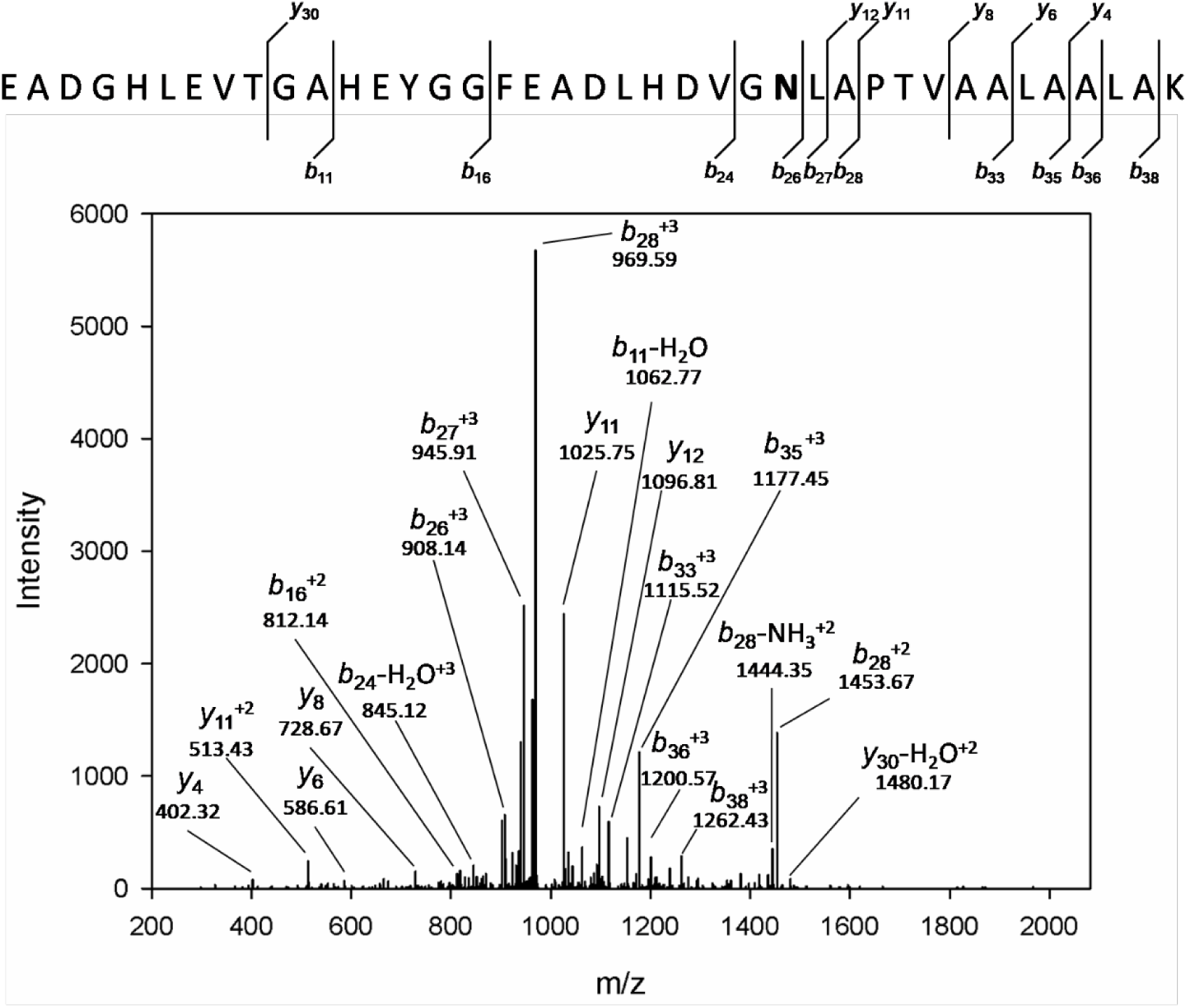
Representative spectra of peptide containing the E321N mutation obtained by LC-MS/MS of *Ms*EPSPS E321N protein. Peptide sequence: EADGHLEVTGAHEYGGFEADLHDVG **N**LAPTVAALAALAK. Point mutation marked in **bold**. Fragment b- and y-ions and their neutral loses are indicated.

**Figure S5.**
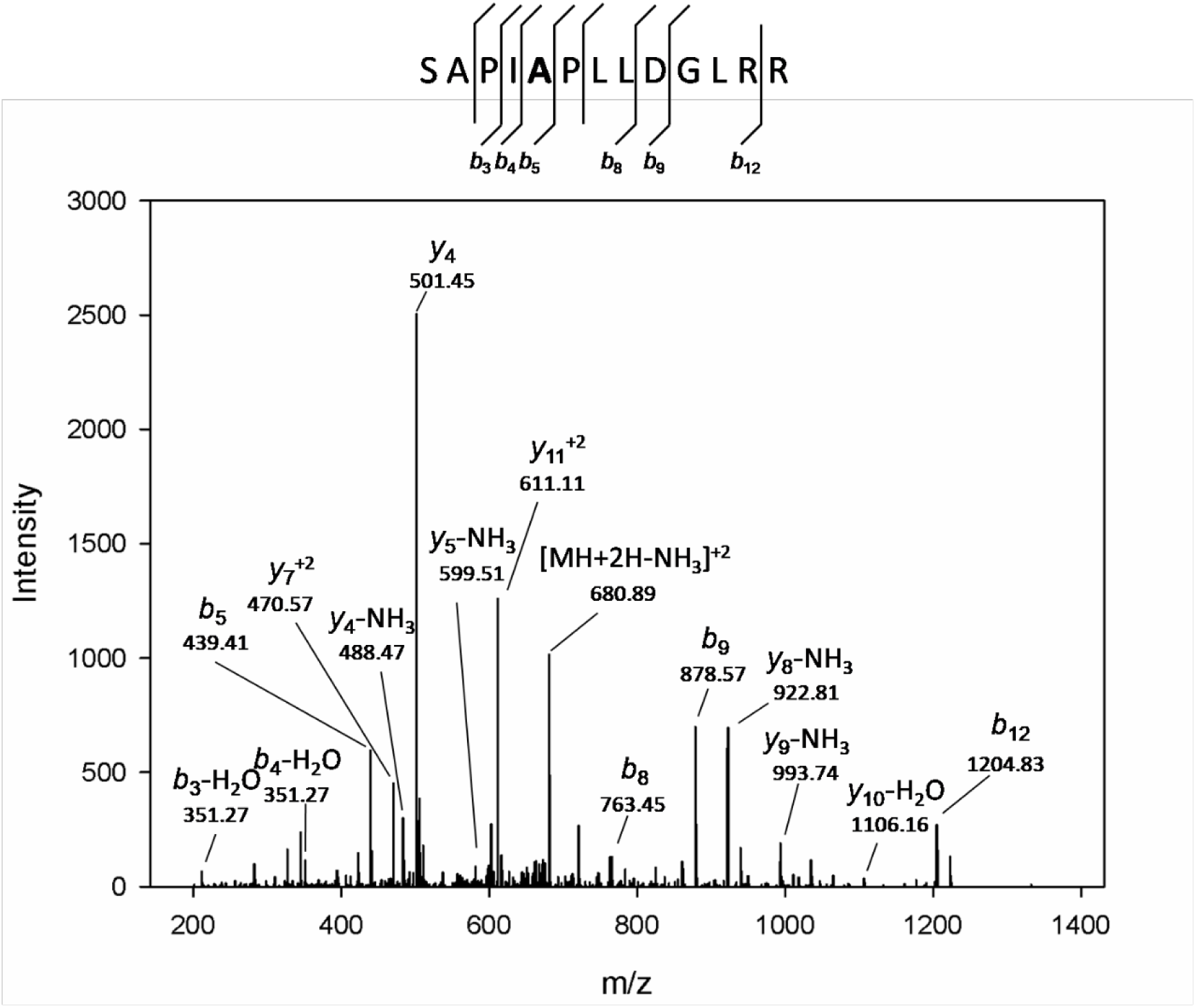
Representative spectra of peptide containing the R134A mutation obtained by LC-MS/MS of *Ms*EPSPS R134A protein. Peptide sequence: S**A**PIAPLLDGLRR. Point mutation marked in **bold**. Fragment b- and y-ions and their neutral loses are indicated.

**Figure S6.**
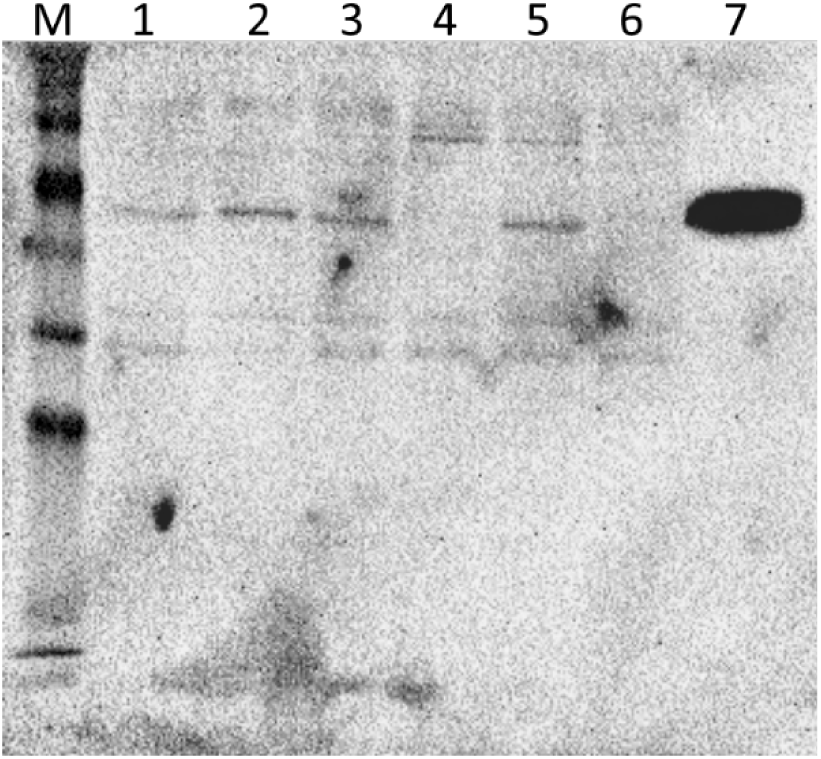
Representative Western Blot from the CRISPRi system, silencing *Ms*EPSPS. Lane M: ProSieve^®^ Color Protein Markers (Lonza). Lane 1-2: PAM1 (1) and PAM3 (2) without induction with ATC (0h). Lanes 3-4: PAM1 18h after induction without (3) and with (4) ATC. Lanes 5-6: PAM3 18h after induction without (5) and with (6) ATC. Lane 7: purified *Ms*EPSPS as positive control.

**Figure S7.**
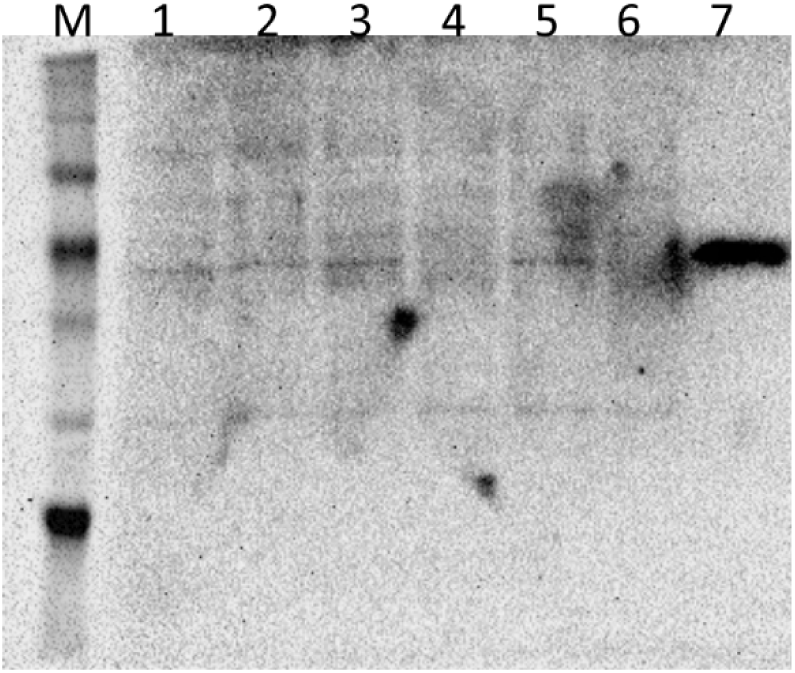
Representative Western Blot from the CRISPRi system, silencing *Ms*EPSPS. Lane M: ProSieve^®^ Color Protein Markers (Lonza). Lane 1-2: duplicate of 0h (without ATC induction) for PAM2. Lanes 3 and 5: duplicate of PAM2 without ATC after 18h. Lanes 4 and 6: duplicate for PAM2 after 18h of induction with ATC. Lanes 7 purified *Ms*EPSPS as positive controls.

**Figure S8.**
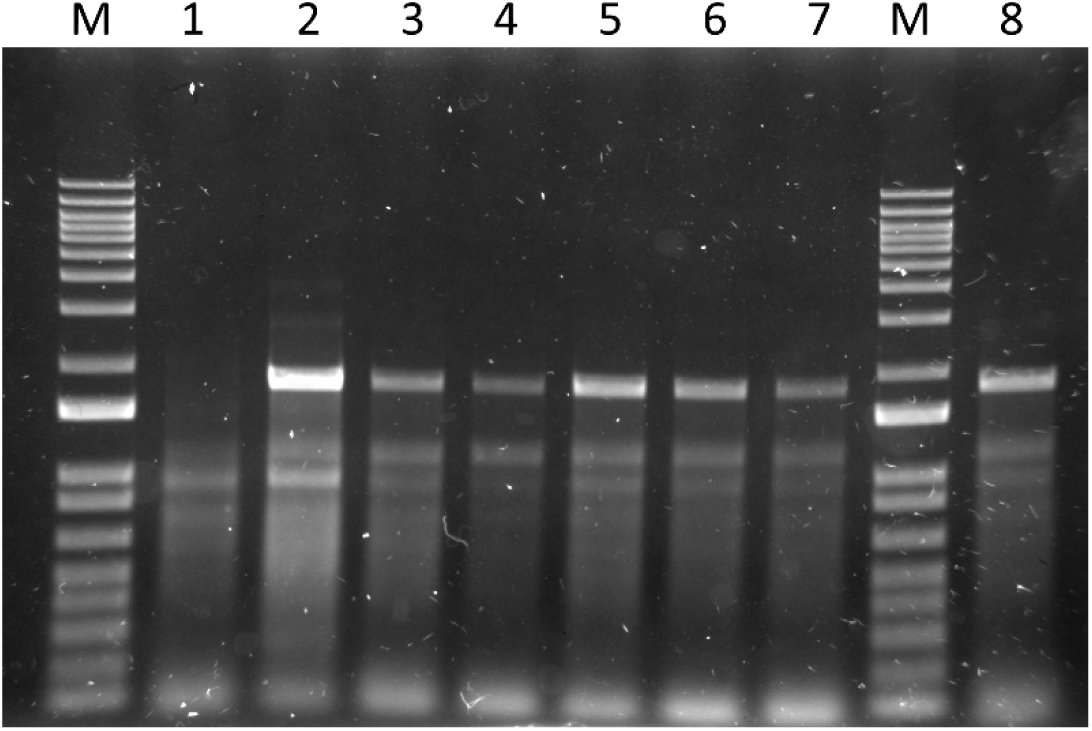
PCR confirming the interruption of the *aroA* gene produced in defined media (7H10), carrying the WT, D61A, or D61W mutant copies. Genomic DNA extracted from white colonies, and a pair of primers specific for the interrupted gene were used for this reaction. A band of 1,813 bp was expected for allelic exchange mutants. Lane M: 1kb plus DNA ladder (Invitrogen). Lane 1: *M. smegmatis* mc²155 genomic DNA (negative control). Lanes 2-6: strains carrying the WT copy of *aroA* gene. Lane 7-8: strains carrying the D61A and D61W mutant copies of *aroA* gene.

